# Identification of frequent activating *HER2* mutations in primary canine pulmonary adenocarcinoma

**DOI:** 10.1101/528182

**Authors:** Gwendolen Lorch, Karthigayini Sivaprakasam, Victoria Zismann, Nieves Perdigones, Tania Contente-Cuomo, Alexandra Nazareno, Salvatore Facista, Shukmei Wong, Kevin Drenner, Winnie S. Liang, Joseph M. Amann, Sara L. Sinicropi-Yao, Michael J. Koenig, Krista La Perle, Timothy G. Whitsett, Muhammed Murtaza, Jeffrey Trent, David P. Carbone, William P. D. Hendricks

## Abstract

Naturally occurring primary canine lung cancers are aggressive malignancies that are increasingly common in pet dogs. They share clinicopathologic features with human lung cancers in never-smokers, but their genetic underpinnings are unknown. Through multi-platform sequencing of 88 primary canine lung tumors or cell lines, we discovered somatic, coding *HER2 (ERRB2)* point mutations in 38% of canine pulmonary adenocarcinomas (cPAC, 28/74), but none in adenosquamous (cPASC, 0/11) or squamous cell (cPSCC, 0/3) carcinomas. In cPASC, *PTEN* was the most frequently mutated gene (18%) while one case each bore likely pathogenic *HRAS, KRAS, EGFR, MET, TP53,* or *VHL* somatic mutations. In cPSCC, no recurrently mutated genes were identified, but individual somatic coding mutations were found in *BRAF* and *PTPN11.* In cPAC, we also identified recurrent somatic mutation of *TP53* (13.5%), *SMAD4* (5.4%), *PTEN* (4.1%), and *VHL* (2.7%). cPACs assessed by exome sequencing displayed a low somatic mutation burden (median 64 point mutations, 19 focal copy number variants, and 1 translocation). The majority (93%) of *HER2* mutations were hotspot V659E transmembrane domain (TMD) mutations comparable to activating mutations at this same site in human cancer. Other *HER2* mutations identified in this study were located in the extracellular domain and TMD. *HER2*^V659E^ was detected in the plasma of 33% (2/6) of dogs with localized *HER2*^V659E^ tumors. *HER2*^V659E^ correlated with constitutive phosphorylation of AKT in cPAC cell lines and *HER2*^V659E^ lines displayed hypersensitivity to the *HER2* inhibitors lapatinib and neratinib relative to *HER2*-wild-type cell lines. These findings have translational and comparative relevance for lung cancer and *HER2* inhibition.

## MAIN TEXT

Naturally occurring primary canine lung cancer is clinically challenging and increasingly common^1^. The disease course and underlying biology resemble that of human lung cancer in never-smokers which, in humans, accounts for 10-25% of lung cancers, causes approximately 26,000 deaths annually, and has a high incidence of erb-B family gene mutations such as those impacting *EGFR*. While incidence of smoking-related lung cancer is decreasing, lung cancer incidence in never-smokers is increasing^2^. Never-smoker lung cancer is primarily the non-small cell histology (NSCLC) arising from lung tissue as opposed to small cell (SCLC) arising in bronchi of smokers. NSCLC histologies include adenocarcinoma (AC) and squamous cell carcinoma (SCC). The etiology of never-smoker lung cancer is also distinct from that of smokers. It is associated with factors including environmental exposures (second-hand smoke, radon, asbestos, arsenic, silica, and pollution) as well as age, sex, family history, and genetic loci^3^. Unique genomic characteristics of human never-smoker lung cancer include low somatic mutation burden, enrichment for C:G>T:A transitions, and somatic activating point mutations or fusions impacting *EGFR* (45%), *ALK* (5-11%), *ROS* (1.5-6%), *HER2* (3-5%), and *RET* (2%)^4^ Five-year overall survival is estimated at 23%, but outcomes are dependent on molecular subtype and treatment regimen. For example, EGFR inhibitors can improve outcomes in EGFR-mutant lung cancers, however 85% of never-smoker lung AC and SCC cases are *EGFR* wild-type (WT) in the United States. Clinical trials of immune checkpoint inhibitors have recently shown improved outcomes for human lung cancers, but analysis of large phase II immunotherapy trials suggests that benefits are limited in low-mutation-burden (≤ 10 mutations/Mb) cases such as those found in smokers^5^. A need exists for improved biologic understanding and development of new models to fuel translational research in never-smoker lung cancer.

Lung cancer in pet dogs has limited standard of care beyond surgery and little is known of its molecular underpinnings^1^. Primary lung tumors typically arise in older dogs (11 years) and resemble human NSCLC histotypes including canine pulmonary adenocarcinoma (cPAC), adenosquamous carcinoma (cPASC), and squamous cell carcinoma (cPSCC). These subtypes collectively represent 13-15% of primary lung tumors^6,7^. Patients are often diagnosed late with lesions incidentally discovered during routine geriatric evaluation or due to nonspecific symptoms including dyspnea (6% to 24%) and cough (52% to 93%) that do not manifest until the tumor is more than 3 cm. These tumors can be diagnostically challenging. Rates at which ultrasound or CT-guided fine needle aspirates of the pulmonary mass provide cytologic diagnosis range from 38% to 90% of cases, varying broadly based on tumor accessibility and aspirate quality. At diagnosis, 71% of malignant canine lung tumors show signs of invasion and 23% show distant metastasis. Partial or complete lung lobectomy is standard of care, dependent on extent of disease spread. Median survival is 345 days for localized disease without nodal involvement where surgical remission can be achieved, but only 60 days when nodes are involved. Responses to cytotoxic chemotherapy (cisplatin, vindesine, doxorubicin, and mitoxantrone) in the setting of disseminated disease are limited. Targeted small molecules and immune checkpoint inhibitors have not been extensively studied in part because the molecular underpinnings of canine lung cancer remain largely unknown. In naturally occurring canine NSCLC, while comprehensive genomic profiling has been limited, *KRAS* hotspot mutation prevalence estimates from targeted studies have varied from 0-25%^7–9^. We have previously shown that EGFR mutation, overexpression, or phosphorylation is rare in cPAC compared to matched non-affected chemotherapy-naive lung tissue whereas significant overexpression and/or phosphorylation of PDGFRα, ALK, and *HER2* are present^10^. We now describe for the first time the detailed genetic underpinnings of primary canine lung cancers through multi-platform next-generation sequencing of 88 cases.

## RESULTS

### The genomic landscape of naturally occurring primary canine lung cancer

In order to map the genomic landscape of primary canine lung cancer we undertook multiplatform next-generation sequencing of 88 NSCLC cases including 77 tumor/normal matched pairs and 11 cell lines (**Table 1**). The cohort included 74 cPAC, 11 cPASC, and 3 cPSCC. Labrador retrievers represented the most commonly affected pure breed dog (21%) with mixed breeds (25%) and multiple other single pure breeds. The predominant cPAC subtype was papillary adenocarcinoma (62%). The cohort was gender-balanced (52% females) and primarily neutered/spayed (92%) with a median age at diagnosis of 11 years. Smoking status in the pet’s household was not available. Extended clinical annotation is shown in **Table S1** and **Figure S1**.

**Table 1.** Genomic analyses performed in primary canine lung cancer.

| Table 1. Genomic analyses performed in primary canine lung cancer |  |  |
| --- | --- | --- |
| Analysis platform | Samples analyzed* | Type of alteration detected |
| Exome | 5 cPAC and matching normal | Germline and somatic SNVs, CNVs, and SVs in coding regions |
| Amplicon | 61 cPAC and matching normal^, 8 cell lines | Germline and somatic SNVs in 51 cancer genes |
|  | 10 cPASC and matching normal, 1 cell line | Germline and somatic SNVs in 51 cancer genes |
|  | 2 cPSCC tumor and matching normal samples, 1 cell line | Germline and somatic SNVs in 51 cancer genes |
| **Total** | **78 tumor-normal pairs, 10 cell lines** |  |

\* an additional cell line (OSUK9PADRiley) was analyzed by sanger sequencing.
^ all matching normal samples were available except one (CCB30381)
cPAC: Canine Pulmonary Adenocarcinoma; cPASC: Canine Pulmonary Adenosquamous Carcinoma;
cPSCC: Canine Pulmonary Squamous Cell Carcinoma.

To identify somatic point mutations, copy number changes, and translocations, we first sequenced the coding genomic regions of five cPAC tumors and matching normal samples using a custom Agilent SureSelect canine exome capture kit comprising 982,789 probes covering 19,459 genes. Tumors were sequenced to mean target coverage of 298X with 99% of target bases covered ≥ 40X (**Table S2**). Constitutional DNA samples were sequenced to mean target coverage of 263X with 99% of target bases covered ≥ 40X. A total of 648 high-confidence somatic SNVs (median 64, range 37-406), 165 focal CNVs (median 19, range 0-116), and 3 SVs (median 1, range 0-1) were identified (**Tables S3-S5** and **Figure 1A, 1B,** and **1C**). We identified non-recurrent, somatic SNVs in genes whose human orthologs have been implicated in human cancer according to COSMIC^11^ Tiers 1 and 2 including nonsynonymous variants in *BCLAF1, CD274, CDH10, CSMD3, FANCD2, FOXP1, KIT, MAML3, NOTCH2, PER1, PTPN13, SEPT9* and *TP53* (**Table S3**). Large-scale recurrent single copy changes (whole chromosome or arm-level gains/losses) included *canis familaris* chromosome (CFA) 1 loss, CFA4 gain, CFA5 loss, CFA11 loss, CFA20 loss, CFA22 gain, CFA25 loss, CFA28 loss, and CFA35 loss (**Figure S2**). Focal CNVs containing COSMIC genes included deletions involving the tumor suppressor *CDKN2A/B* locus (40% of cases) and *PTEN* (20%), as well as amplifications involving the oncogenes *AKT1* (20%), *KIT*(20%), *KDR* (20%), and *TERT*(20%) (**Figure 1B and Table S4**).

**Figure 1.**
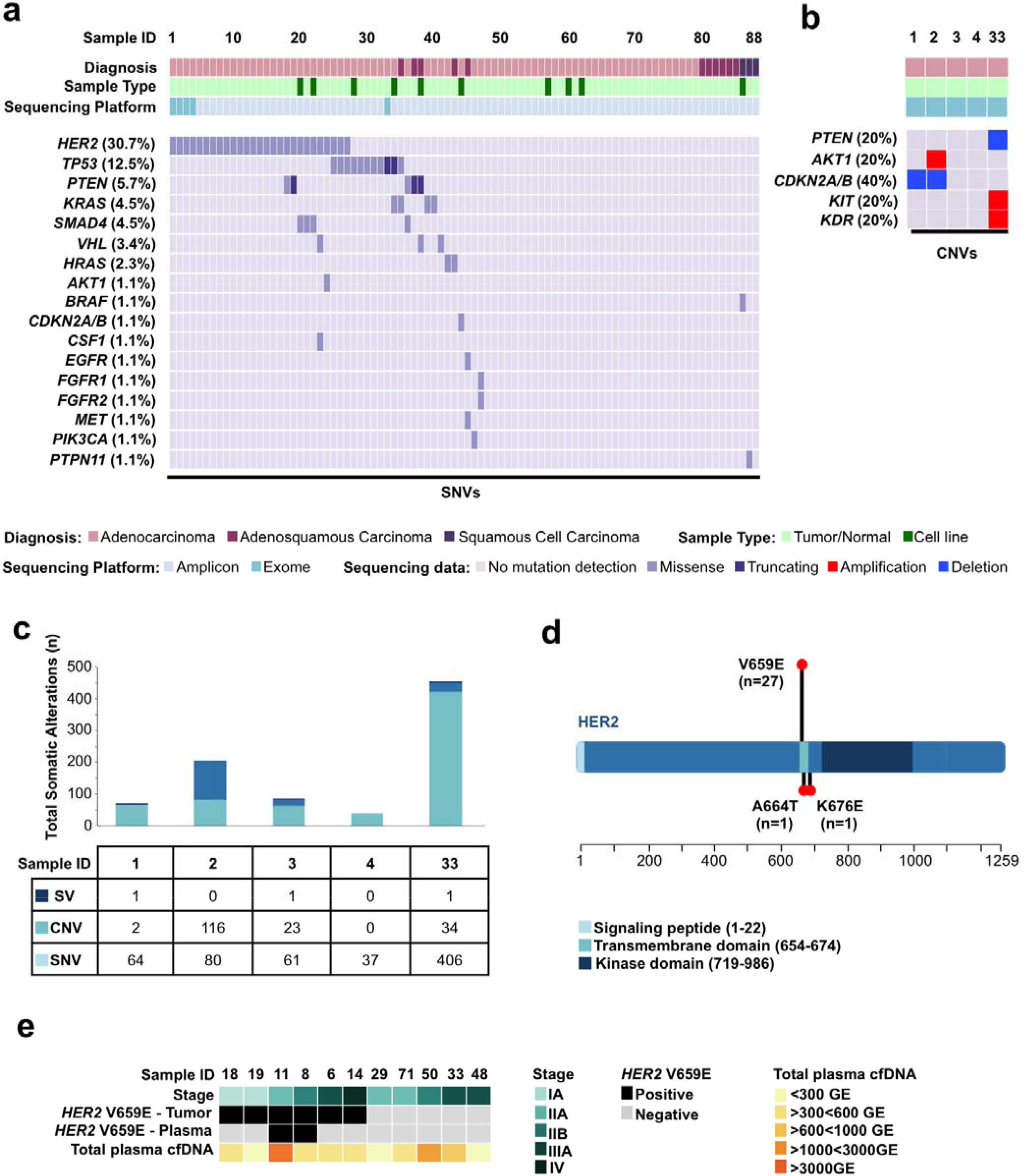
The Genomic Landscape of Primary Canine Lung Cancers. **(A)** Recurrent likely pathogenic somatic mutations in cancer genes identified in primary canine lung cancers through multi-platform sequencing. SNVs were determined from combined tumor/normal exome and/or amplicon sequencing across 88 total tumors and cell lines. **(B)** CNVs were determined from tumor/normal exome data in five cPAC cases. **(C)** Somatic mutation burden (SNVs, CNVs, and SVs) identified by exome sequencing of five tumor/normal cPAC cases. **(D)** Distribution of somatic *HER2* mutations within the *HER2* protein identified in primary canine lung cancers. The length of the lollipops is proportional to the number of mutations found. **(E)** Detection of *HER2* hotspot mutations in plasma from 11 canine primary lung cancer cases.

Of the three somatic SVs, only the *WWTR1-ATP5F1* translocation impacted a COSMIC gene, *WWTR1.* The sole gene bearing recurrent nonsynonymous SNVs was *HER2* (80%) with the missense mutation V659E occurring in three cases and the missense mutation K676E in a fourth case (**Figure 1D**). No *HER2* amplifications were detected in these five tumors. We also assessed somatic point mutation signatures according to their trinucleotide context (**Figure S3**)^12,13^. The most common signature in these five cases was the age-associated COSMIC Signature 1A in 4/5 (80%) featuring frequent C>T substitutions at NpCpG trinucleotides. COSMIC Signature 2, associated with APOBEC cytidine deaminase activity, was also present in two cases, CCB050227 and OSU431895.

To identify somatic point mutations across a broader cohort of canine lung cancers, we used a custom canine cancer amplicon-based next generation sequencing panel to assess 281 coding regions (**Table S6**) of canine orthologs of 57 frequently mutated human cancer genes in 73 additional lung tumors (61 cPAC, 10 cPASC, 2 cPSCC), two previously exome-sequenced tumors with matched normal tissue, and 10 cell lines (8 cPAC, 1 cPASC, and 1 cPSCC). These cases were sequenced to an average depth of 3,383x (**Table S7**). A median of 1 somatic coding point mutation (range 0-3) within sequenced panel regions was identified across all cases. Likely pathogenic recurrent point mutations included *HER2* V659E (29.8%), *KRAS* G12D/V (3.4%), *SMAD4* D351Y/G (3.4%) and *TP53* R239Q/G (2.2%) (**Table S8**). Two additional somatic mutations in *HER2* including A664T and K676E were identified in single cases (**Figure 1D**). Overall, recurrently mutated genes containing somatic potentially pathogenic SNVs included *TP53* (12.5%), *PTEN* (5.7%), *SMAD4* (4.5%), *KRAS* (4.5%), *VHL* (3.4%) and *HRAS* (2.3%). Finally, based on both exome and amplicon sequencing, we evaluated germline SNPs to identify putatively pathogenic rare variants (i.e. those not previously identified in dogs based on review of presence in dbSNP 151^14^ and/or ≥10% frequency in DogSD^15^) in 81 genes previously reported to potentially be associated with susceptibility to lung cancer in humans^16^. Based on these criteria, we identified nine rare putatively pathogenic SNPs in five dogs in the genes *CHRNA3, CYP1B1, DNAH11* and *HER2* (**Table S9**). Of these SNPs, the only variant with an equivalent in its human orthologous gene was *DNAH11* R1460W corresponding to human *DNAH11* R1444W (rs1035326227, MAF < 0.01%), but the human SNP has not been associated with a disease phenotype. The *HER2* variants, both V1189I, occurred in two cases for which no somatic *HER2* mutations were identified in tumor tissue. The human orthologous position, V1184, has not been shown to bear variation in human data. Though rare, the canine variant has been identified in 4% of cases in DogSD and, based on functional effect prediction (FATHMM), it is likely neutral. Finally, none of the genes bearing rare SNPs showed second somatic hits in tumor tissue and thus, overall, data to support their pathogenicity is limited.

We additionally performed matched tumor/normal amplicon sequencing to evaluate the genomic landscapes of 11 cPASC and 3 cPSCCs, subtypes which are under-studied entities in dogs and humans, especially in non-smokers (**Figure 1A and Table S8**). In contrast with cPAC, no *HER2* mutations were identified in these tumors. In cPASC, *HRAS* Q61L and *KRAS* Q61K each occurred in one case. Thus 18% of cases bore *RAS* hotspot mutations. *PTEN* stop gains additionally occurred in 2/11 (18%) of cases at high tumor allele frequencies and were exclusive with *RAS* mutations. Additional likely pathogenic somatic mutations also occurred in a single cancer gene in a single tumor each including *EGFR* A726T, *MET* M1269V, *TP53* R147C, and *VHL* P97L. Finally, while no recurrent mutations were identified in the three cPSCCs, we identified one case with a somatic *BRAF* V588E and another bearing *PTPN11* G503V.

### *HER2* is frequently mutated in canine pulmonary adenocarcinoma (cPAC)

*HER2* was the most frequently mutated gene in our multi-platform next generation sequencing cohort, occurring exclusively in cPACs (27/74, 36.5%) with two mutations occurring in a single patient (**Figure 1A**). We additionally identified a *HER2* mutation in the cell line OSUK9PAPADRiley solely by Sanger sequencing of the codon 659 locus. We thus identified 29 total *HER2* mutations overall (**Figure 1D**). In 24 cases, the *HER2* variants were evaluated on at least two platforms including exome sequencing, amplicon sequencing, Sanger sequencing, and/or droplet digital PCR (ddPCR, **Table S10**). The *HER2* variant tumor allele fraction (AF) median by amplicon sequencing was 21.3% (range 8.4 – 51.9%). All low AF (< 20%) cases identified by amplicon sequencing were also validated by Sanger and/or ddPCR. Some low AF cases were not able to be visualized on Sanger traces, but were detectable by ddPCR. Notably, one cell line, OSUK9PAPADOscar, contained a low AF *HER2* V659E variant (AFs of 15% by amplicon and 16% by ddPCR) during early short-term culture (passage 4) that was no longer detectable by Sanger sequencing or ddPCR in later passages (passage 15). Importantly, passage 15 was utilized for all functional studies described below and it was thus considered *HER2*^WT^ in this setting. Overall, V659E missense mutations located in the *HER2* TMD occurred in 93.3% of *HER2*-mutant cases. Additional *HER2* mutations included A664T (OSU419040) and K676E (CCB050354), which have not been previously described in orthologous human *HER2* regions. *HER2* mutations co-occurred with mutations in *TP53, SMAD4, VHL, PTEN, AKT1* or *KDR. TP53* was found in 3/27 (11%) of the 27 *HER2*-mutant tumors, although it was more commonly mutated in *HER2* wild-type tumors. Three of the four SMAD4 mutations in this cohort were comutated in *HER2*-mutant tumors (11%). Additional co-occurring mutations included *PTEN* in 2/27 (11%) and *VHL, AKT* and *KDR* each co-mutated in a single *HER2*-mutant case.

### *HER2* mutations are detectable in canine plasma DNA

Cell-free tumor DNA (ctDNA) in plasma has been increasingly used for noninvasive genotyping in human cancer patients^17^. To develop a canine blood test that could rapidly identify dogs with *HER2*-mutant lung cancer, we investigated whether cPAC *HER2* hotspot mutations are detectable in ctDNA. We evaluated plasma from 11 dogs, 5 with *HER2*-wild-type tumors and 6 with *HER2*^V659E^ tumors using droplet digital PCR (ddPCR) and custom primers targeting 57 bp surrounding codon 659 (**Table S11**). We analyzed wild-type tumor samples, plasma DNA from unrelated commercially available canine plasma samples and template-free controls to establish assay specificity (BioreclamationIVT #BGLPLEDTA-100-P-m). Using uniform gating for all experiments, we found 1/7 template-free samples showed 1 wild-type droplet and no samples showed any evidence of mutant DNA amplification. In wild-type tumor and plasma DNA samples, 2/8 samples showed 1 mutant droplet each. Based on these results, we required at least three mutant droplets to confidently detect *HER2*^V659E^. We detected *HER2*^V659E^ in 6/6 positive control tumor DNA samples where we had previously identified V659E mutations using amplicon sequencing. In these samples, we observed a high correlation between allele fractions measured using amplicon sequencing and ddPCR (Pearson’s r 0.976, p=0.0008). In 11 plasma samples from dogs with cPAC tested using ddPCR, median total cell-free DNA yield was 1.6 (range 0.3-10.0) (**Table S11**). Requiring at least three mutant droplets to support mutation detection, the median limit of detection (LoD) for mutation allele fraction was calculated at 0.61% (range 0.10%-3.11%). *HER2*^V659E^ mutations were detected in 2/6 plasma samples from dogs with *HER2*^V659E^-positive tumors at 1.9% and 2.3% allele fractions. Both ctDNA-positive cases had stage II cancer, consistent with low levels of ctDNA observed. *HER2*^V659E^ was not detected in any plasma samples from dogs with *HER2*^WT^ tumors, confirming assay specificity (**Figure 1E**).

### HER2 expression in primary canine lung cancer

In human cancers, *HER2* bears activating point mutations, focal amplifications or copy number gains, and overexpression (by qRT-PCR and/or immunohistochemistry). Amplification and overexpression are mutually exclusive with point mutations. To assess *HER2* copy number gains and overexpression, *HER2* copy number was first determined in the five exome-sequenced cases (**Figure S2** and **Table S4**). No large-scale CFA9 gains or focal *HER2* amplifications were detected. However, these cases predominantly bore somatic putatively activating point mutations and might not be expected to contain concomitant gains. Therefore, we evaluated *HER2* RNA and protein expression by qRT-PCR and IHC. RNA samples from 49 lung tumors within this cohort (9 *HER2*-mutant) alongside 14 normal lung tissue samples distal to tumor areas but from the same lung lobe were evaluated. Median *HER2* expression fold-change relative to expression of the house-keeping gene HPRT in normal lung samples was 1.06 (range 0.28-4.11) and in tumors was 0.85 (range 0.07-4.50) (**Figure S4**). No significant difference in relative *HER2* expression was observed between tumor and normal or *HER2*-mutant and *HER2*^WT^ groups.

Additionally, in order to quantify *HER2* protein expression in cPAC, digital image analysis was performed on eight tumors from FFPE. Three of the samples bore the *HER2*^V659E^ hotspot mutations, one bore *HER2*^A664T^, and four were *HER2*^WT^ according to amplicon and Sanger sequencing. All cases were positive for *HER2* staining with homogeneous and diffuse staining of tumor cell cytoplasm and cell membrane, but no staining in adjacent stroma or vasculature (**Figure S5**). Positive staining was observed in bronchial epithelium of the adjacent non-affected lung in all cases. Consistent with absence of observed *HER2* amplifications, no significant differences (mean ± SEM) were detected in the tumor positivity percentage for *HER2* (47±5.4 and 35±5.1) between the WT and *HER2*-mutant groups, respectively. No significant differences in *HER2* staining were present for percent minimum (51±2.9 WT *vs.* 55±5.1, *HER2* mutations) or percent maximum (97±0.31 WT *vs.* 96±0.47 *HER2* mutations) stain intensity (**Table S12**). Overall, most tumors showed moderate expression of *HER2* based on qRT-PCR and IHC with some variability, but levels were typically consistent with those seen in normal tissue and did not vary based on *HER2* mutation status.

### HER2 is constitutively active in *HER2*^V659E^ cPAC cell lines

In human cancers, *HER2* V659E mutations increase *HER2* autophosphorylation, EGFR phosphorylation, and activation of phosphatidyl-inositol-3-kinase (PI3K), and mitogen-activated-protein-kinase (MAPK) pro-survival signaling pathway members (e.g. AKT and ERK) relative to wild-type *HER2*^18^. To determine whether *HER2*^V659E^ constitutively activates downstream signaling in cPAC, we first validated *HER2* genotype in seven canine lung cancer cell lines through amplicon sequencing, ddPCR or Sanger sequencing of the V659 locus, Sanger sequencing of all *HER2* coding regions, and/or aCGH to determine *HER2* copy number status as previously published^19^ (**Table S10**). One cell line, OSUK9PAPADOscar, bore a low allele frequency *HER2*^V659E^ mutation when sequenced by amplicon panel at low passage (passage 4) as a primary culture, but had lost this allele in later established passages (passage 15) characterized by Sanger sequencing and ddPCR and utilized for functional studies. We thus evaluated *HER2* activation in one *HER2*^V659E^ cPAC cell line, OSUK9PAPADRex, and three *HER2*^WT^ cell lines (two cPAC – CLAC and OSUK9PAPAPADOscar – and one cPASC – OSUK9PADSQ) by Western blotting for total and phospho-AKT in the presence and absence of the ErbB ligand, neuregulin (hNRG) post-serum starvation. Only the *HER2*^V659E^ line, OSUK9PAPADRex, showed constitutively high AKT phosphorylation post-starvation even in the absence of hNRG stimulation (**Figure 2A**).

**Figure 2.**
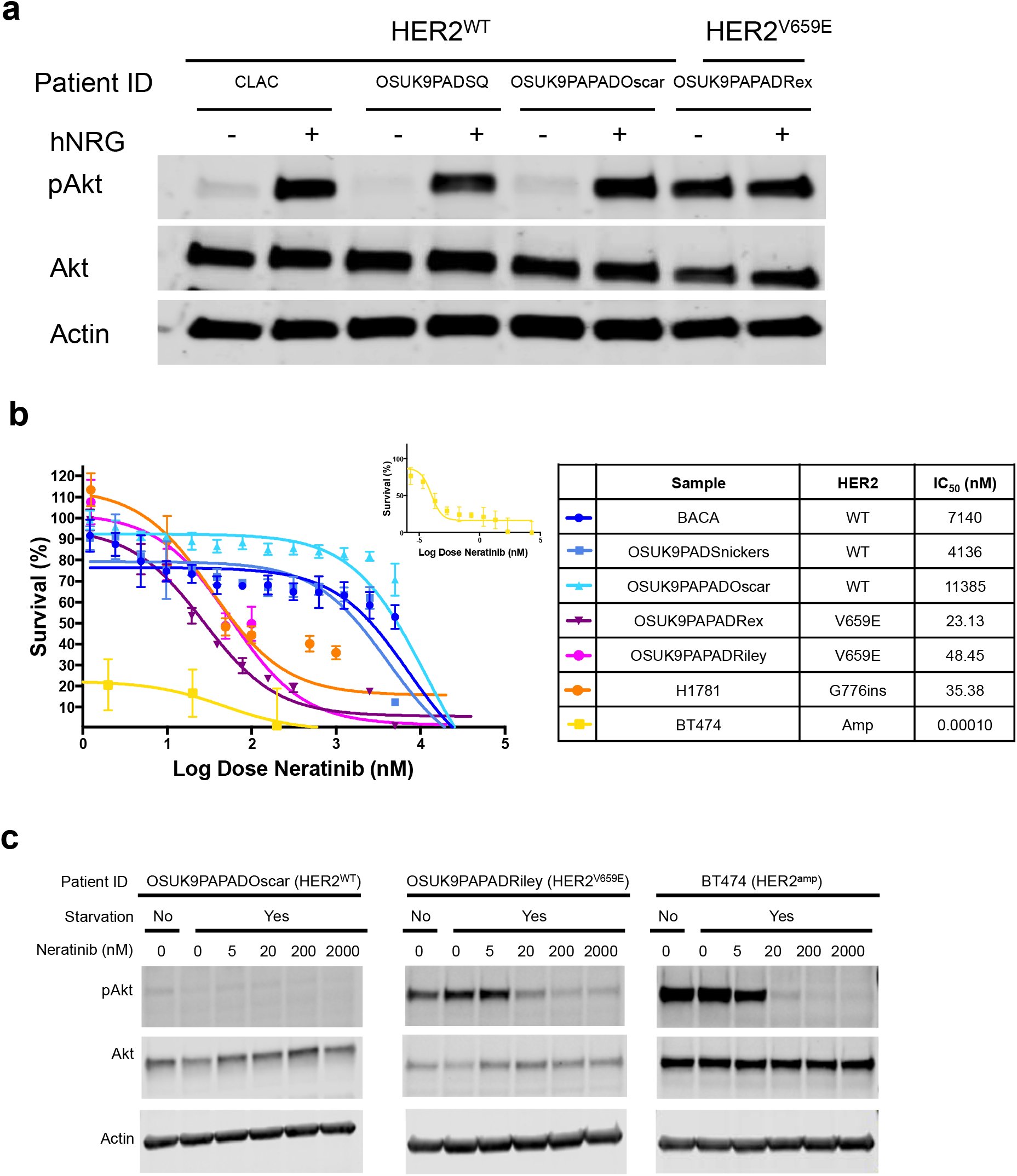
*HER2*^V659E^ constitutively activates downstream *HER2* signaling and is associated with response to *HER2* inhibitors in primary canine pulmonary adenocarcinoma (cPAC) cell lines. **(A)** *HER2* signaling activation in canine lung cancer cell lines. Phospho-AKT and AKT levels were assessed by Western blot under serum starvation in the presence and absence of EGFR activation by hNRG in *HER2*^V659E^ and *HER2*^WT^ cPAC cell lines. **(B)** Neratinib drug-dose-response studies in primary lung cancer cell lines. Five canine cell lines (three *HER2*^WT^ and two *HER2*^V659E^) and two human lung cancer positive controls with known *HER2* activating mutations (BT474 – *HER2*-amplified, and H1781 – *HER2*^G776V^) and *HER2* inhibitor responses were treated with 14 neratinib doses ranging from 100 μM to 5.5×10^−2^ nM for 72 hours with CellTiterGlo viability endpoints. Survival is shown relative to DMSO vehicle control.

### *HER2*^V659E^ cPAC cell lines are selectively sensitive to HER2 inhibitors

*HER2*^V659E^ has been shown to stabilize receptor dimerization and constitutively activate downstream pro-survival signaling^20^. Therefore, anti-*HER2* drugs may prove effective for selectively eradicating *HER2*^V659E^ tumors. We evaluated differential sensitivity of *HER2*^V659E^ and *HER2*^WT^ canine lung cancer cell lines to the small molecule *HER2* inhibitors lapatinib and neratinib. Five cPAC cell lines (two *HER2*^V659E^ and three *HER2*^WT^) and two *HER2*-mutant human cancer cell lines – BT474 (*HER2*^AMP^) and H1781 (kinase domain *HER2*^G776ins^), both of which are known to respond to *HER2* inhibitors^18,21^ – were treated with neratinib and four (one *HER2*^V659E^ and three *HER2*^WT^) with lapatinib for 72 hours (**Figures 2B and S6**). Significant differences in viability were observed between *HER2*^V659E^ and *HER2*^WT^ cPAC cell lines for both drugs with IC_50_s < 200 nM in *HER2*^V659E^ versus IC_50_s > 2500 nM in *HER2*^WT^. All *HER2*-mutant cell lines were sensitive to neratinib with IC_50_s < 50 nM (**Figure 2B**) significantly lower than *HER2*^WT^ responses (p=0.0079). We additionally observed a neratinib dose-dependent decrease in p-AKT in the *HER2*-mutant cell lines OSUK9PAPADRiley (*HER2*^V659E^) and BT474 (*HER2*^amp^) whereas p-AKT levels in OSUK9PAPADOscar (*HER2*^WT^) were low at all treatment levels (**Figure 2C**). These studies further support that *HER2*^V659E^ in cPAC is an activating event that confers dependency on downstream signaling and sensitivity to targeted *HER2* inhibition.

## DISCUSSION

Through multi-platform next-generation sequencing of 88 naturally occurring primary canine NSCLC cases (77 tumors and 11 cell lines), we describe for the first time the detailed genomic underpinnings of this cancer. The cohort included major NSCLC subtypes occurring in dogs and humans: cPAC (n=74, cPASC (n=11), and cPSCC (n=3) (**Table S1, Figure S1**). Although lung cancer may be over-represented in Doberman pinschers, Australian shepherds, Irish setters, and Bernese mountain dogs^7^, Labrador retrievers comprised the largest pure breed in this cohort (21%) followed by mixed breeds (25%) and multiple other pure breeds in smaller numbers. The cohort was gender-balanced (52% females), primarily neutered/spayed (92%), and bore a median age at diagnosis of 11 years. Although second-hand smoke exposure in these dogs remains possible given that exposure was not recorded, genomic landscapes of human lung cancers in never-smokers have not been shown to differ based on exposure to second-hand smoke^22^. Exposure to other environmental carcinogens such as air pollutants may also play a role in development of lung cancers. For example, increased lung cancer risk may be present in dogs with higher amounts of carbon deposits known as anthracosis^23^, although in humans anthracosis has been commonly observed in normal lungs as well as tumors and lymph nodes. A mutational etiology for such pollutant exposure has also not been established. In this cohort, anthracosis was recorded in 15 cases and pneumoconiosis (lung disease associated with pollutant exposure) in one case. However, no associations between anthracosis annotation and genetic features of these cases were observed. Overall, our studies included broad representation of lung cancer across histologic subtypes, breeds, ages, and pollutant exposures reflective of primary canine lung cancer diversity seen in the clinical setting in the United States.

Unique genomic characteristics of human never-smoker lung cancer include low somatic mutation burden, C:G>T:A enrichment, and activating mutations or fusions impacting *EGFR* (45%), *ALK* (5-11%), *ROS* (1.5-6%), *HER2* (3-5%), and RET(2%)^4,24^. Here, we also observed a low somatic burden of SNVs, CNVs, and SVs through exome sequencing in five matched tumor/normal cPAC pairs. We additionally observed that the most common mutation signature in these five cases was the age-associated COSMIC Signature 1A in 4/5 (80%) featuring frequent C>T substitutions at NpCpG trinucleotides and similar to the C>T substitution enrichment seen in human NSCLC (**Figure S3**). This signature is associated with age in many human cancers, putatively the result of spontaneous deamination of 5-methyl-cytosine. COSMIC Signature 2, associated with APOBEC cytidine deaminase activity, was also present in two cases. This signature, characterized by C>T and C>G substitutions at TpCpN, is most prominently associated with cervical and bladder cancers, but is also commonly found in 16 cancer types including lung adenocarcinoma and squamous cell carcinoma. While these signatures are sometimes associated with *APOBEC* gene variants in human cancers^25^, no putatively pathogenic germline or somatic *APOBEC* mutations were observed. Based on our studies, primary canine lung cancers bear a low mutation burden and mutation signatures reflective of those seen in human never-smoker lung cancers.

The most common recurrently mutated genes containing somatic potentially pathogenic SNVs in the full cohort included *HER2* (31.5%), *TP53* (12.5%), *PTEN* (5.7%), *SMAD4* (4.5%), *KRAS* (4.5%), *VHL* (3.4%) and *HRAS*(2.3%). Recurrent *CDKN2A/B* focal deletions were also observed in 2/5 (40%) cases (**Figure 1A, 1B**). While *HER2* contained the largest number of somatic mutations identified in this study, the broader driver mutation landscape provides an informative view of the genetic underpinnings of primary canine lung cancer. *CDKN2A* deletions were the most common alteration by frequency overall, occurring at rates comparable to those in human NSCLC. Two focal deletions were observed out of five exome-sequenced cases with signs of larger-scale CFA11 losses in remaining cases (**Figure S2**). We additionally identified a single case bearing a homozygous missense mutation, G50R, equivalent to codon G101 mutations in human cancer (14 reports according to COSMIC). *CDKN2A* is mutated in ~30% of all human NSCLC, primarily via homozygous deletion, and this number is reduced to around 25% in never-smokers. The next most common alterations after *CDKN2A* and *HER2* were *TP53* missense and truncating mutations comparable to DNA binding domain mutations in human *TP53.* Similar to human NSCLC, we observed a reduced burden of *TP53* mutations (12.4%) relative to human smoker NSCLC in which more than half of tumors are mutated. We found two stop gains and nine missense mutations. Notable likely pathogenic missense mutations with human equivalents include R164H (human R175, 1,857 mutations in COSMIC), R239Q/G (human R249, 739 mutations in COSMIC), and R272H (human R282, 955 mutations in COSMIC).

*PTEN*mutations were the next most common alterations at 5.6%. *PTEN* is mutated in ~9% of human NSCLC, but only ~2% of never-smoker NSCLC. In addition to a homozygous deletion in a single exome-sequenced case, *PTEN* mutations were predominantly stop gains in three of five cases bearing somatic SNVs. The remaining two *PTEN* mutations were missense, occurring in the C2 domain and included D230Y (comparable to human pathogenic D252Y) and C228R (no comparable human variant). We additionally identified four somatic mutations in the tumor suppressor *SMAD4,* mutated in ~5% of human NSCLC at comparable rates in smoker and never-smoker cancer. These mutations included three D351 codon mutations and W524C, all in the MH2 domain, comparable to those recurrently described in human lung cancer. The SMAD4 protein shares 99% identity between dog and human with D351 being the 4^th^ most commonly mutated codon (45 mutations in human lung cancer from COSMIC).

*KRAS* mutations are the most common oncogenic mutations in human smoker NSCLC (~30-40% of cases), but occur at reduced frequencies in never-smoker lung cancer (0-7%). We observed a low frequency of *KRAS* mutations. The KRAS protein shares 100% identity between dog and human with the G12 codon in the nucleotide binding domain being the most commonly mutated region. *KRAS* mutations in our cohort (2 G12V, 1 G12D, and 1 Q61K) were comparable to human hotspots in codons G12 and Q61. Missense mutations in canine *HRAS* were also located in human-equivalent hotspots (Q61L, F78S). Additional likely pathogenic somatic mutations included individual cases of *AKT1* amplification, *KIT/KDR* amplification, *EGFR* A726T (human A755), *MET* M1269V (human M1268), and *VHL* P97L (human P97). *WWTR1,* the only COSMIC gene bearing a somatic translocation in exome-sequenced cases, has been shown to undergo translocation with *CAMTA1* in human epithelioid hemangioendothelioma^26^. Here, we identified a *WWTR1* translocation of unknown consequence with *ATP5F1.* Overall, although we observed a similar mutation spectrum relative to human never-smoker NSCLC, a notable exception is lack of *EGFR* mutations which, although present in few smoker lung cancers (0-7%), are enriched in human never-smokers (~45%). Further, although we identified translocations occurring in coding regions in five exome-sequenced tumors, it remains possible that, as in human never-smoker lung cancer, *EML4-ALK* fusions, *ROS1* fusions, *RET* fusions, and other fusions may be present in canine lung cancer.

In addition to charting the landscape of cPAC, we have found recurrent *KRAS* and *TP53* mutations in cPASC and provide a view of possible drivers in cPSCC. In cPASC, *HRAS* Q61L and *KRAS* Q61K each occurred in one case (18% of cases thus with *RAS* mutations. Finally, while no recurrent mutations were identified in the three cPSCCs, we identified one case with somatic *BRAF* V588E (equivalent to the human V600E hotspot) and another bearing *PTPN11* G503V (equivalent to the human G503V hotspot).

*HER2* contained the largest number of somatic mutations with high frequency hotspot mutations occurring solely in cPAC (37.8%). *HER2* is a well-characterized human oncogene and drug target mutated in ~6% of all cancers based on cBioPortal query of 10,967 cases in the TCGA pan-cancer atlas^27,28^. The majority of these alterations are focal amplifications, but activating point mutations are also common. In human NSCLC, *HER2* mutations have been identified as oncogenic drivers in ~1-4% of cases, mostly in exon 20 at codon 776 resulting in constitutive *HER2* kinase domain activation and downstream signaling through PI3K and MAPK pathways^24,29,30^. It may also be more commonly mutated in human never-smoker lung cancer, with point mutations at frequencies of 3-5%^31^. *HER2*-mutant human NSCLC is predominately female never-smokers with adenocarcinomas who carry a median OS of ~2 years^30^. *HER2* TMD polar mutations (*HER2*^V659E/D^, *HER2*^G660D^) such as those we have discovered in cPAC are present in 0.18% of human lung adenocarcinomas and are exclusive with *HER2* kinase domain mutations^32^. We determined *HER2* status in canine lung cancer through multi-platform analysis. Our amplicon analysis covered canine *HER2* exons 8 and 17-22 including transmembrane and kinase domains. Additionally, we performed Sanger sequencing of all exons in five canine cell lines with wild-type *HER2* based on amplicon sequencing (OSUK9PAD, BACA, CLAC, K9PADSQ and OSULSCC1) and found no somatic *HER2* mutations in other sites (**Table S10**). It is nonetheless possible that somatic mutations occurring in other regions of *HER2* were not identified in amplicon-sequenced samples even though data facilitating functional interpretation of these variants would be limited.

In addition to point mutations, *HER2* amplification has also been identified in ~1% of human NSCLC^24^, with enrichment in EGFR-inhibitor-resistant tumors^33^. Protein overexpression has been reported in 6-35% of tumors including up to 42% of adenocarcinomas and has been shown to correlate with poor prognosis^34–37^. We detected no somatic *HER2* focal amplifications or large-scale CFA9 gains in five exome-sequenced cases (**Figure 1B** and **S2**) or two previously aCGH-profiled cell lines. However, four of these seven cases contained somatic, putatively activating *HER2* SNVs. Given that *HER2* amplification/overexpression and SNVs are typically mutually exclusive, it remains possible that our broader amplicon cohort contained undetected *HER2* amplifications. We therefore utilized qRT-PCR and IHC studies to more broadly assess *HER2* overexpression and did not find evidence for significant tumor-specific *HER2* overexpression (**Figures S4** and **S6** and **Table S12**). Thus, it is unlikely that *HER2* is frequently amplified in primary canine lung cancer. Rather, point mutation appears to be the primary mechanism of *HER2* hyperactivation.

We have additionally shown that *HER2* hotspot mutations can be detected in the plasma of dogs bearing *HER2*^V659E^ cPACs even at early stages of the disease (at least stage II) (**Figure 1E** and **Table S11**). In human NSCLC, ctDNA has been shown to be significantly enriched in plasma relative to controls with key genetic features identifiable via liquid biopsy. Associations have been found between ctDNA levels and tumor stage, grade, lymph node status, metastatic sites, response, and survival^38,39^. In fact, the first FDA-approved liquid biopsy test for lung cancer was the cobas EGFR Mutation Test v2, a real-time PCR assay utilized in NSCLC for the detection of *EGFR* exon 18-21 mutations in tissue or plasma to guide EGFR inhibitor treatment assignment^40,41^. Our proof-of-principle study supports that ctDNA is also detectable in primary canine lung cancer patient plasma. A non-invasive *HER2*^V659E^ assay will enable genotyping patients when tumor tissue is limited and may have a role in treatment monitoring or detection of minimal residual disease. This assay will also facilitate prospective analysis of *HER2*^V659E^’s diagnostic and prognostic value.

Canine *HER2* shares normal and oncogenic roles with human *HER2* based on sequence conservation and prior study of its role in cell signaling in the dog. It shares 92.2% amino acid identity with human *HER2*. *HER2*^V659E^ occurs at a highly conserved residue identical to the human variant (100% identity with human *HER2* in the TMD from amino acids 654-674) and to the *neu* (rat *HER2*) variant identified in a rat glioblastoma cell line that originally led to discovery of *HER2’s* oncogene status^42^. *HER2* has been implicated in canine cancers via overexpression by IHC and qRT-PCR in canine mammary tumors^43^, through its utility as a vaccine target in canine osteosarcoma^44^, and through downstream signaling hyperactivation in canine lung cancer^10^. In human cancers, *HER2* TMD polar mutations like those discovered here, have been shown to constitutively activate pro-survival *HER2* signaling^32^ and have been associated with responses to *HER2* inhibitors^20^. We have confirmed in this study that, similar to human *HER2* TMD mutants, canine *HER2*^V659E^ cell lines constitutively activate downstream signaling through AKT and are selectively sensitive to *HER2* inhibitors *in vitro* including lapatinib and neratinib (**Figure 2** and **S6**).

We have charted the genomic landscape of primary canine lung cancers including the NSCLC subtypes cPAC, cPASC, and cPSCC. We have identified recurrent *HER2* mutations in these cancers and present, to our knowledge, the first complete suite of evidence supporting an oncogenic role for and dependency on constitutively activating mutations in *HER2* in a canine cancer. Further work is needed to more comprehensively profile these tumors, particularly according to variation across breeds, and to determine the prognostic significance of the mutations described here. However, these data offer immediate diagnostic and therapeutic opportunities for dogs with primary lung cancer. They aid comparative understanding of never-smoker and *HER2*-mutant lung cancer across species.

## ONLINE METHODS

### Sample Collection

Tumors and cell lines from 89 dogs from the Canine Comparative Oncology and Genomics Consortium (CCOGC)^45^ and The Ohio State University (OSU) College of Veterinary Medicine Biospecimen Repository were included. Veterinary pathologists board certified by the American College of Veterinary Pathologists (ACVP) confirmed tumor diagnosis based on histopathology. This study was approved by The OSU Institutional Animal Care Use Committee (2010A0015-R2). Tumor and normal tissue samples were flash frozen in liquid nitrogen or formalin-fixed and paraffin-embedded (FFPE). Cell lines were maintained in RPMI 1640 with GlutaMAX™ (Gibco™, Thermo Fisher Scientific #61870036) supplemented with 10% heat inactivated fetal bovine serum at 37°C and 5% CO_2_ and passaged at 90% confluence. Cell lines with known passage data (except BACA) were sequenced within the first eight passages of derivation and subsequently expanded for phenotypic studies. Canine cell lines were authenticated by IDEXX BioResearch (Columbia, MO) using Cell Check Canine STR Profile and Interspecies Contamination Tests on January 30, 2018 or by NkX2 (or TTF-1) RT-PCR. Human cell lines included BT474 (ATCC #HTB20, *HER2* focal amplification) and H1781 (ATCC #CRL-5894, *HER2*^G776V^). Whole blood was processed and stored within one hour of collection. Blood for cell-free DNA and germline DNA extraction was collected in 10ml K2 EDTA Blood tubes (ThermoFisher Scientific #22-253-145). Plasma separation was performed at room temperature within 1h (2x serial centrifugation at 2000 rpm × 15 min). Plasma aliquots were stored frozen at – 80°C. DNA extraction from plasma, white blood cells and tissue was performed with MagMAX Cell-Free DNA Isolation Kit (Thermo Fisher Scientific #A29319), DNeasy Blood and Tissue Kit (QIAGEN #69504) and Qiagen AllPrep DNA/RNA Mini Kit (QIAGEN #80204), respectively.

### Exome Sequencing and Analysis

Informatic tools, versions, and flags are shown in **Table S13**. We utilized a custom Agilent SureSelect canine exome capture kit with 982,789 probes covering 19,459 genes. Exome libraries were sequenced on the Illumina HiSeq2000 producing paired end reads of 85bp. FASTQ files were aligned to the canine genome (CanFam3.1) using BWA v0.7.8. Aligned BAM files were realigned and recalibrated using GATK v3.3.0 and duplicate pairs were marked with Picard v1.128 (http://broadinstitute.github.io/picard). Somatic copy number variants (CNVs), and structural variants (SVs) were called with tCoNutT (https://github.com/tgen/tCoNuT) and DELLY v0.76 respectively. Somatic single nucleotide variants (SNV) were identified from two or more of the following callers: Seurat v2.6, Strelka v1.0.13 and MuTect v1.1.4. Germline SNVs were called using Haplotype Caller (GATK v3.3.0), Freebayes and samtools-Mpileup. Variant annotation was performed with SnpEff v3.5. The SomaticSignatures R package was used to identify mutation signatures.

### Targeted Amplicon Sequencing and Analysis

We developed a custom canine cancer amplicon sequencing panel consisting of 281 amplicons targeting exons and hotspot regions in 57 genes, with amplicon sizes ranging from 91-271 bp (**Table S6**). We pooled primers in two multiplexed pools to separate adjacent amplicons and any amplicons that showed cross-amplification using *in silico* PCR. We prepared sequencing libraries using digital PCR amplification following the manufacturer’s protocols for the ThunderBolts Cancer Panel (RainDance Technologies) with modifications as previously described^46^. Sequencing was performed on the Illumina MiSeq generating paired-end 275 bp reads. Sequencing reads were demultiplexed and extracted using Picardtools. Sequencing adapters were trimmed using ea-utils and fastq files were assessed for quality using FASTQC.

Sequencing reads were aligned to CanFam3.1 using bwamem-MEM^47^. Custom in-house scripts based on samtools were used to create pileups for every sample. Pileups were analyzed in R to call SNVs and indels. For each potential non-reference allele at each targeted locus in a sample, we evaluated the distribution of background noise across all other sequenced samples. To call a variant, we required the observed non-reference allele is an outlier from the background distribution with a Z-score > 5. In addition, we required tumor depth ≥100x, allele frequency ≥10%, number of reads supporting the variation ≥10, and allele fraction in the germline sample <1%. Finally, variant calls were manually curated by visualization in IGV v2.3.71. All next generation sequencing data (exome and amplicon) has been deposited in NCBI repository under accession number PRJNA523624.

### Sanger Sequencing

23 primer pairs covering all exons of *HER2* were designed using Primer 3 (http://bioinfo.ut.ee/primer3-0.4.0/) including a universal M13 tag. Amplicons were Sanger sequenced at the DNASU sequencing facility at Arizona State University on an ABI 3730XL (Applied Biosystems, Foster City, CA) and analyzed with Mutation Surveyor DNA Variant Analysis Software (SoftGenetics, State College, PA).

### HER2 Inhibitor Drug Dose-Response Studies

*HER2* inhibitors lapatinib (Selleckchem, #S2111) and neratinib (Puma Biotechnology, Los Angeles, CA), were assessed in 10-16 point 72 hr drug-dose response screens (from 2×10^−7^ nM to 100 μM) with CellTiter-Glo^®^ luminescent cell viability assay (Promega, #G7570). Cells were cultured in RPMI supplemented with 10% FBS and 1% Penicillin/Streptomycin. Luminescence was read using Synergy Mx (Biotek) plate reader. Six replicates were performed for each dose. Curve-fitting and IC_50_ calculations were performed using GraphPad Prism v7.00 (GraphPad Software, La Jolla, CA).

### Droplet Digital PCR

*HER2*^V659E^ genotyping was performed on tumor samples and plasma cell-free DNA with droplet digital PCR (ddPCR). 50 μl reactions contained 2X KAPA PROBE FAST Master Mix (Kapa Biosystems, #KK4701), 200 μM dNTP Mix (New England BioLabs Inc, #N0477S), 1x Droplet Stabilizer (RainDance Technologies, #30-00826), 1 μM pooled primer mix (IDT), 1 μM mutant (FAM labeled) or wild type (TET labeled) *HER2* V659E probe (IDT), and 21.75 μL of template DNA. Each reaction was partitioned into 10 million droplets using the RainDrop Digital PCR Source instrument (RainDance). PCR amplification was performed as follows: 1 cycle 3 min at 95 °C, 50 cycles 15 sec at 95 °C and 1 min at 60 °C with a 0.5 °C/sec ramping from 95 °C to 60 °C, 1 cycle 98 °C for 10 min and hold at 4 °C. Droplet fluorescence was measured using RainDrop Digital PCR Sense instrument and analyzed using accompanying RainDrop Analyst II Software v.1.0 (RainDance). Primer and probe sequences used for *HER2*^V659E^ detection in ctDNA were Forward: 5’-CCCACGACCACAGCCA-3’, Reverse: 5’-CCCTGTGACATCCATCATTGC-3’ and Probe: 5’-CAGAATGCCC(T/A)CCACAGC-3’.

### qRT-PCR

cDNA was obtained by reverse transcription with iScript (Biorad, #1708891) and samples were subjected to *HER2* (target) and HPRT1 (reference) amplification in a QuantStudio™ 6 Flex Real-Time PCR System under standard conditions with Syber Green technology (KapaBiosystems, #KK4602). Primer sequences were: *HER2*-Forward: 5’-CATCTGCACCATTGATGTCTA-3’, *HER2*-Reverse: 5 ‘-GGCCCAAGTCTTCATTCTGA-3’, HPRT1-Forward: 5’GCAGCCCCAGCGTCGTGATT-3’, HPRT1-Reverse: 5’CATCTCGAGCAAGCCGCTCAGT-3’. Data was analyzed with Quantstudio Real Time PCR software v1.1. Values for ΔCt, ΔΔCt, and fold changes were calculated as follows: ΔCt=Ct *HER2* – Ct HPRT1; ΔΔCt=ΔCt tumor sample–ΔCt average of normal samples; and fold change =2^(-ΔΔCt).

### Immunohistochemistry

*HER2* protein expression was evaluated on FFPE sections (4 μm) of normal lung and tumor mounted on SuperFrost™ Plus glass slides (Fisher Scientific, #12-550-15). Slides were deparaffinized in xylene and rehydrated in an ethanol gradient. Antigen retrieval was performed with 1mM EDTA adjusted to pH 9.0. An autostainer (Dako, model S3400) was used to carry out immunostaining. A *HER2* rabbit monoclonal antibody that detects the amino terminus of human *HER2*/ErbB2 (Cell Signaling Technology, #4290) was used at 1:400 in antibody diluent with background reducing components (Dako, #S3022) for 30 min. Slides were rinsed, incubated with secondary biotinylated rabbit anti-goat IgG (Vector Laboratories, BA-1000) diluted 1:200 in protein block for 30 min, rinsed, then incubated for 30 min in an avidin/biotin peroxidase system for detection of biotinylated antibodies (VECTASTAIN^®^ Elite^®^ ABC System, #PK-6100). IHC positive controls for *HER2* tyrosine kinase receptor expression were single tissue samples of two canine complex mammary carcinomas^48^. Negative controls were performed on all tissues using a universal rabbit negative isotype control not directed against any known antigen (Dako, #IR600).

### Quantitative Image Analysis

Immunostained and control 1×3-inch microscope slides were scanned at 40X on a high-resolution Scanscope XT (Leica Biosystems) at The Ohio State University Comparative Pathology & Mouse Phenotyping Shared Resource. For quantification of immunoreactivity, images were imported into Visiopharm Image Analysis software (Visiopharm, Hørsholm, Denmark version 2017.27.0.3313), segmented into areas of tumor, necrosis, and normal lung tissue using color labels for each tissue type. *HER2* connectivity was scored using the modified 10007 – *HER2*, Breast Cancer APP (Visiopharm). Thresholds were adjusted to match specimen *HER2* stain intensities for accurate scoring. Area (μm^2^) was quantified for each tissue type and percentages derived from specimen total tissue area. Tumor areas were further segmented into staining and non-staining categories. Percentages of stained and non-stained tumor were calculated based on total tumor area in μm^2^. Maximum, mean, and minimum intensities were also quantified using a built-in software calculation. Staining is expressed as percentage of stain present with 100% equal to black (maximum dark brown) and 0% equal to white (no stain present). Initial thresholds and tissue types were established and mark-ups reviewed in consultation with a pathologist board-certified by the American College of Veterinary Pathologists to ensure accurate measurements and to differentiate between tissue types.

### Immunoblot Analyses

Subconfluent cells were serum starved overnight, then treated with 20 nM neuregulin for 15 min prior to harvest. Cells were lysed in RIPA buffer (50 mM Tris, pH 8.0, 150 mM NaCl, 1% IGEPAL CA-630, 0.5% deoxycholate, 0.1% SDS) with cOmplete™ Mini Protease Inhibitor (Roche, #11836153001) and PhosSTOP™ (Roche, # 4906845001) and loaded in Laemmli buffer at 1 μg/μl. Samples were heated to 95°C for 5 min, then 25 μg were separated on 4-15% SDS-PAGE Criterion Gels (BioRad, #5671085) and transferred to Immobilon-FL PVDF membrane (MilliporeSigma, #IEVH7850). Membranes were dried for 1 hour to overnight, rehydrated in methanol, and washed. Membranes were blocked for 1 hour in LiCor blocking buffer (1% fish gelatin, 1x Tris Buffered Saline, pH 7.4, 0.02% sodium azide). Antibodies were diluted in primary antibody buffer (1% fish gelatin, 1x Tris Buffered Saline, pH 7.4, 0.1% Tween-20, 0. 02% sodium azide) and added to the membranes for overnight incubation at 4°C. After washing, membranes were incubated for 1 h at room temperature with fluorescence-conjugated secondary antibodies conjugated in secondary antibody buffer (1% fish gelatin, 1x Tris Buffered Saline, pH7.4, 0.1 % Tween-20, 0.02% SDS, 0.02% sodium azide). After washing, membranes were scanned using the LiCor Odyssey CLx instrument. Primary antibodies were AKT (CST #4691S, 1:1000), phospho-AKT (CST #4060P, 1:1000), and β-actin (CST #4970S, 1:1000).

## ACKNOWLEDGEMENTS

This study was supported by National Canine Cancer Foundation GL150H-005, the Bisgrove Scholars Program, philanthropic support to the TGen Foundation, CTSA UL1TR001070, NCI P30 CA016058, Brooke’s Blossoming Hope, and the PetCo Foundation. A portion of the tissue samples were provided by the Pfizer Canine Comparative Oncology and Genomics Consortium (CCOGC) Biospecimen Repository. Other investigators may have received specimens from the same subjects in the CCOGC. We thank Dr. Amy LeBlanc, Director of the NCI’s Comparative Oncology Program (COP), and Christina Mazcko, COP Program Manager, for their assistance with CCOGC sample procurement and annotation. Additionally, we thank Dr. Holly Borghese of the OSU CVM Veterinary Biospecimen Repository for her expertise in sample procurement and Dr. Kurtis Yearsley of the OSU Pathology Imaging Core for his expertise in quantitative image analysis. We thank Drs. Alshad Lalani, Irmina Diala, and Lisa Eli of Puma Biotechnology for fruitful discussion of *HER2* inhibition and provision of neratinib for this study.

## AUTHOR CONTRIBUTIONS

Study conception and design: G.L, V.Z, J.T, D.P.C and W.H; Data acquisition: G.L, V.Z, T.C, S.F, S.W, W.S.L and N.P; Analysis and interpretation of data: G.L, K.S, V.Z, N.P, M.M, T.C, AN, S.F, S.W, W.S.L, J.M.A, S.L.S, M.J.K, K.L.P, T.G.W, J.T, D.P.C, W.H; Drafting of manuscript: G.L, K.S, V.Z, N.P, W.H; Critical revision: G.L, K.S, V.Z, J.T, D.P.C, W.H.

## SUPPLEMENTAL TABLES AND FIGURES

Table S1. Extended clinical and multiplatform annotation of primary canine lung cancer cases.

Table S2. Sequencing metrics for primary canine lung cancer exome analysis.

Table S3. Somatic coding SNVs identified by exome sequencing of primary canine lung cancers.

Table S4. Somatic coding CNVs identified by exome sequencing of primary canine lung cancers.

Table S5. Somatic coding SVs identified by exome sequencing of primary canine lung cancers.

Table S6. Canine genomic regions covered by custom amplicon panel.

Table S7. Sequencing metrics for primary canine lung cancer amplicon analysis.

Table S8. Somatic coding SNVs identified by panel sequencing of primary canine lung cancers.

Table S9. Germline SNPs in COSMIC Tier 1 cancer genes identified by exome and panel sequencing in primary canine lung cancers.

Table S10. Multi-platform validation of *HER2* mutations.

Table S11. Non-invasive detection of *HER2*^V659E^ in the plasma of primary canine lung cancer patients.

Table S12. *HER2* protein expression and quantification by immunohistochemistry.

Table S13. Informatic tools utilized in primary canine lung cancer genomic analyses.

**Figure S1. Extended clinical annotation of primary canine lung cancer patients**. Cohort distribution of: (A) Affected breeds, (B) Age at diagnosis, (C) Primary tumor location distribution, (D) Sex, (E) Adenocarcinoma subtype, and (F) Treatment. Yr=years.

**Figure S2. Somatic copy number plots derived from exome sequencing of five primary canine pulmonary adenocarcinomas (cPAC) and matched constitutional DNA**. Tumor copy number states determined by tCoNutT analysis of tumors and matched constitutional DNA from five cPAC cases is shown with each canine chromosome plotted on the x-axis (shown in alternating green and black) and log_2_ fold change shown on the y-axis.

**Figure S3. Somatic mutation signatures identified by exome sequencing in primary canine lung cancers**. (A) The distribution of somatic single nucleotide mutation types in their trinucleotide context from tumor/normal exome sequencing of five cPAC cases. (B) The most common mutation signatures based on trinucleotide context and frequency of somatic single nucleotide mutations from tumor/normal exome sequencing of five cPAC cases. COSMIC Signature 1A (C>T substitutions at NpCpG trinucleotides that are associated with age) was present in four cases. COSMIC Signature 2 (C>T and C>G substitutions at TpCpN, associated with APOBEC cytidine deaminase activity) was present in two cases.

**Figure S4. *HER2* expression in primary canine lung cancer based on quantitative real-time PCR analyses**. Box plots are shown for *HER2* 2^(-ΔΔ Ct) expression fold-change relative to the housekeeping gene HPRT (x-axis) in 49 tumors and 14 matched normal lung tissues comparing (A) Normal lung tissue versus tumor tissue, and (B) Normal lung tissue versus *HER2*-wild-type tumor tissue versus *HER2*-mutant tumor tissue.

**Figure S5. *HER2* cellular location and function in primary canine lung cancer**. (A) Canine papillary adenocarcinoma with intense, complete, circumferential membrane (white arrow) and lateral cytoplasmic membrane (black arrow) anti-*HER2* antibody positive staining (brown) in a patient with wild-type *HER2*. (B) Canine papillary adenocarcinoma with moderate cytoplasmic (black arrow) anti-*HER2* antibody positive staining (light brown) in a patient with wild-type *HER2*. x 40; bar 50 μm. (C) Anti-*HER2* immunohistochemistry of a Grade 1 canine papillary adenocarcinoma wild type for *HER2*. x 20. (D) Segmentation mark-up of the tumor from adjacent normal lung. Tumor is identified by green, whereas red is area within tumor that contains no tissue, and yellow represents areas of non-tumor such as necrosis or tumor stroma. x20.

**Figure S6. Canine primary lung cancer cell line sensitivity to lapatinib**. Four canine cell lines (three *HER2*^WT^ and one *HER2*^V659E^) were treated with 14 lapatinib doses ranging from 100 μM to 5.5×10^−2^ nM for 72 hours with CellTiterGlo viability endpoints were measured and shown as percent survival relative to DMSO vehicle control.

