## Supplementary figures and images for "Identification of frequent activating *HER2* mutations in primary canine pulmonary adenocarcinoma"

### Supplemental Figure 1

#### Slide 1
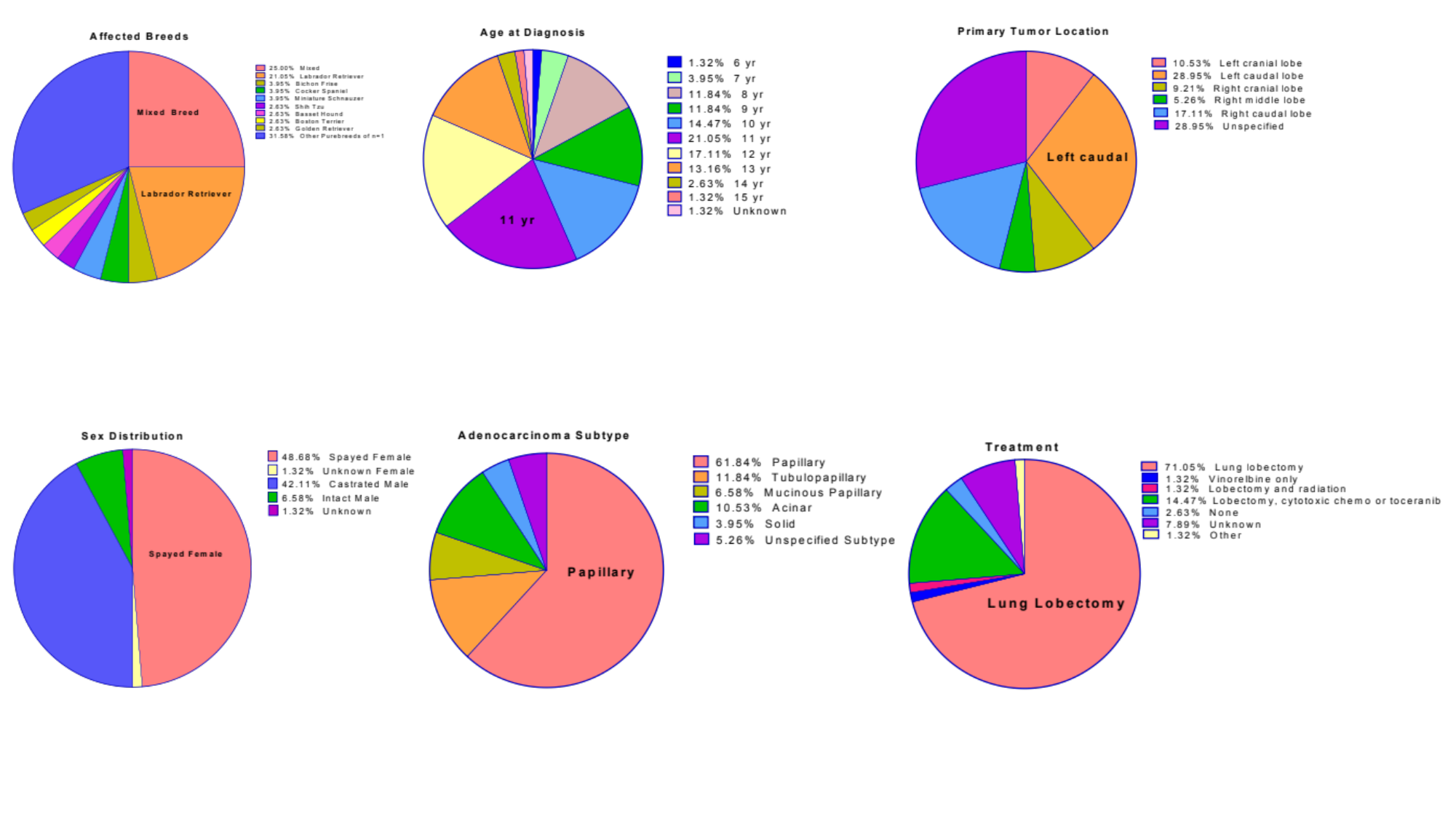

### Supplemental Figure 2

#### Slide 1
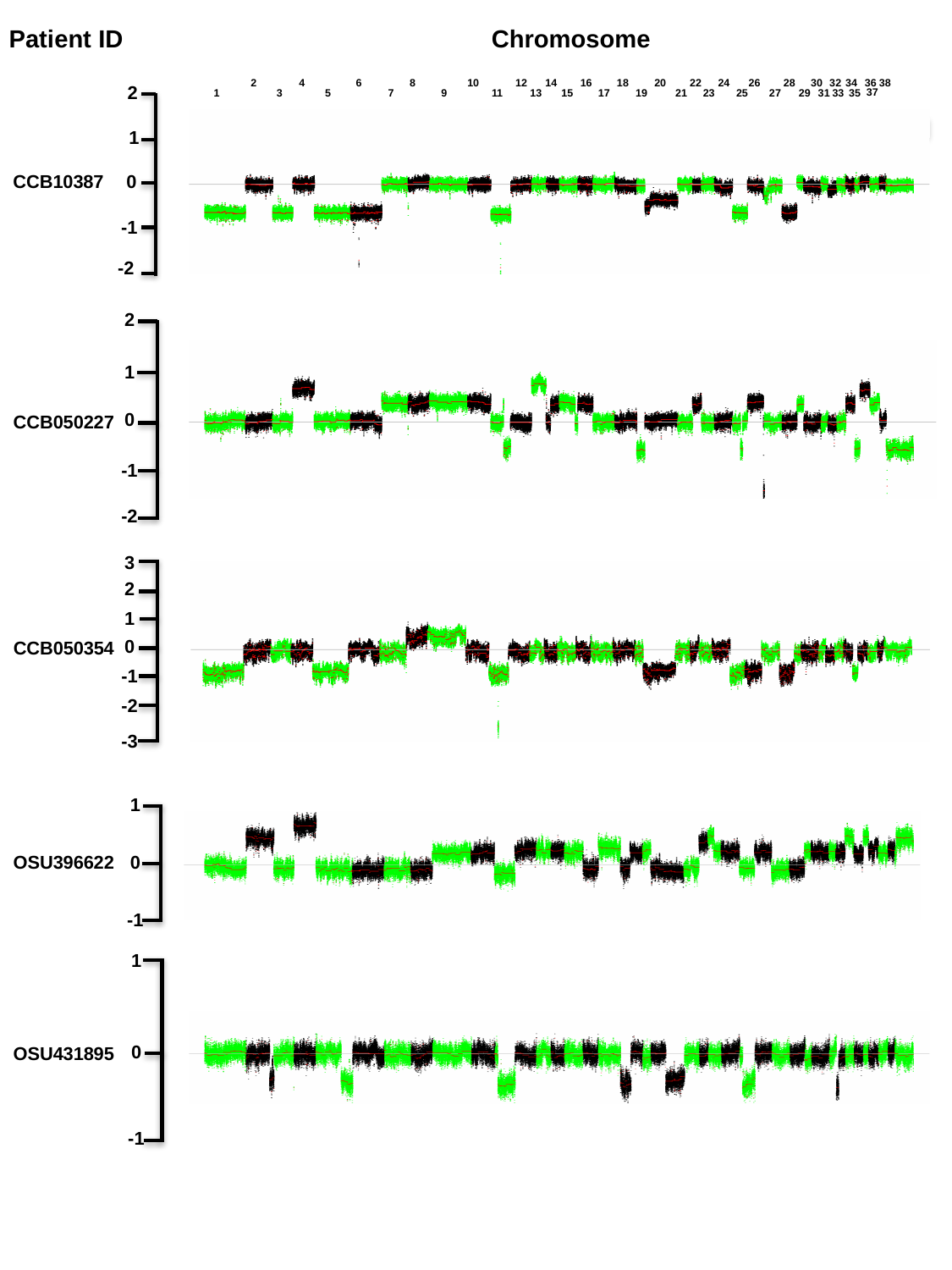

Patient ID
Chromosome
2
4
6
8
10
12
14
16
18
20
22
24
26
28
30
32
34
36
38
2
37
1
3
5
7
9
11
13
15
17
19
21
23
25
27
29
31
33
35
1
CCB10387
0
-1
-2
2
1
0
CCB050227
-1
-2
3
2
1
0
CCB050354
-1
-2
-3
1
0
OSU396622
-1
1
0
OSU431895
-1

### Supplemental Figure 5

#### Slide 1
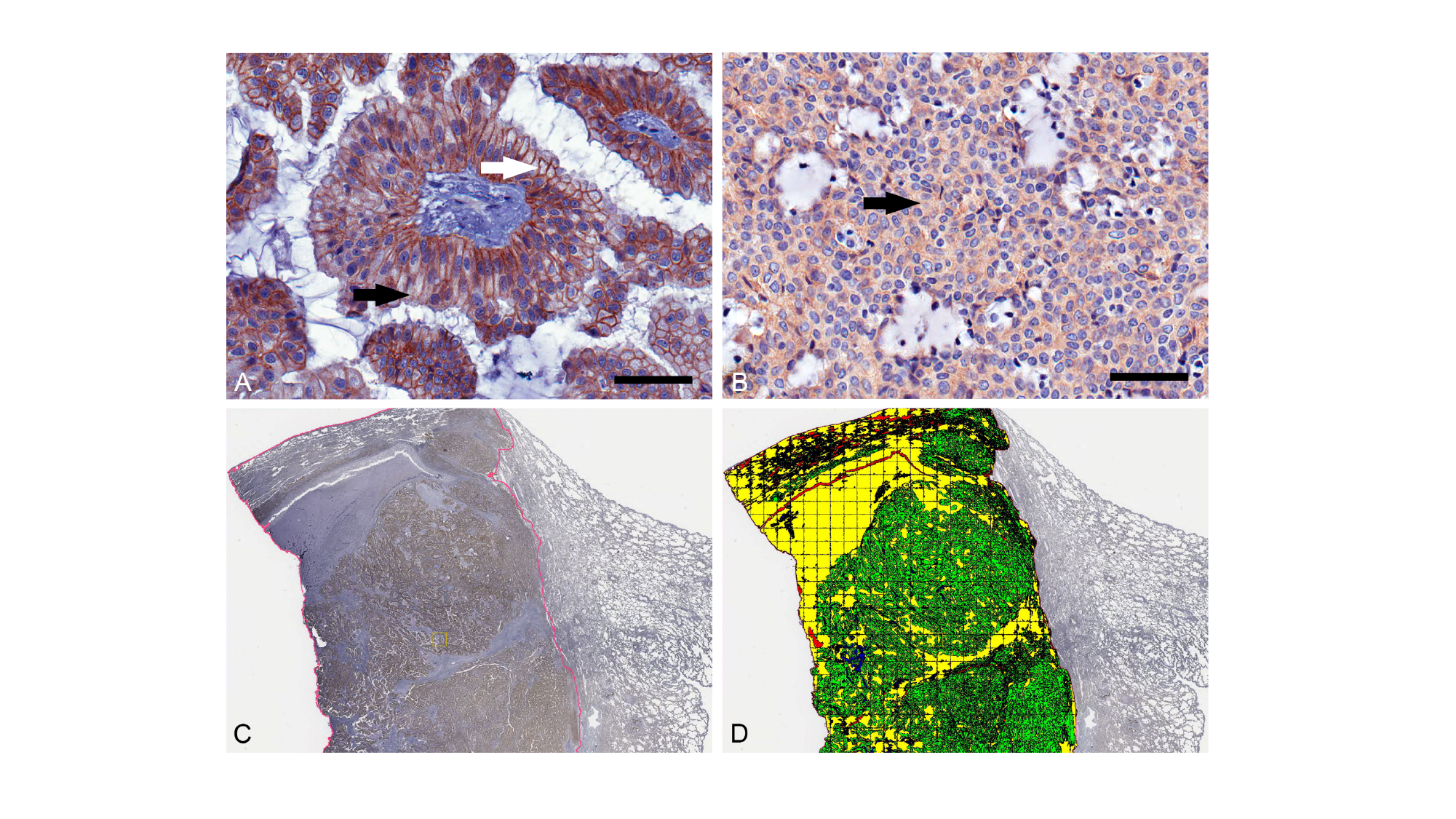

### Supplemental Figure 6

#### Slide 1
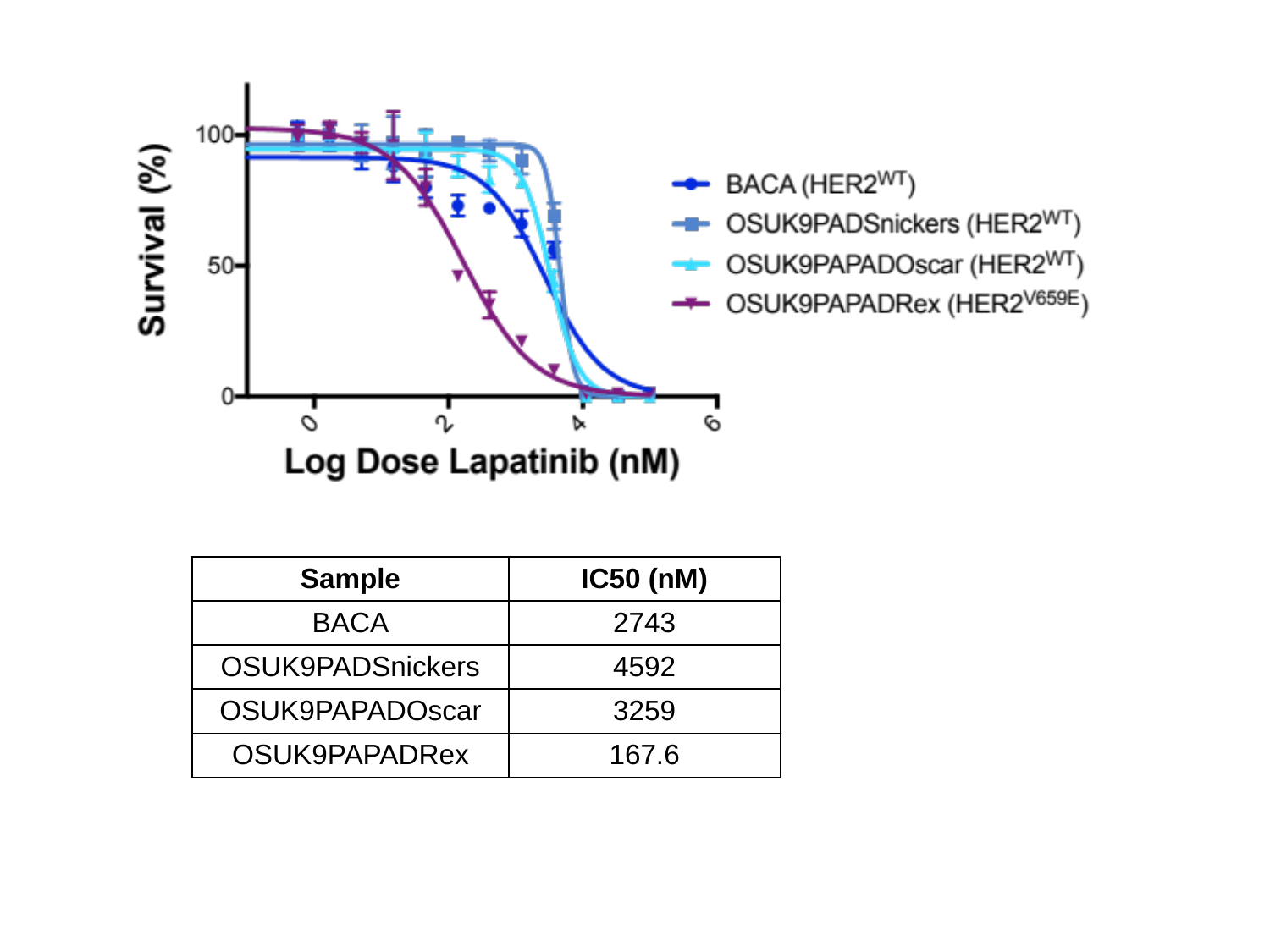

| Sample | IC50 (nM) |
| --- | --- |
| BACA | 2743 |
| OSUK9PADSnickers | 4592 |
| OSUK9PAPADOscar | 3259 |
| OSUK9PAPADRex | 167.6 |
