## Supplemental Figure 3 for "Identification of frequent activating *HER2* mutations in primary canine pulmonary adenocarcinoma"

### Slide 1
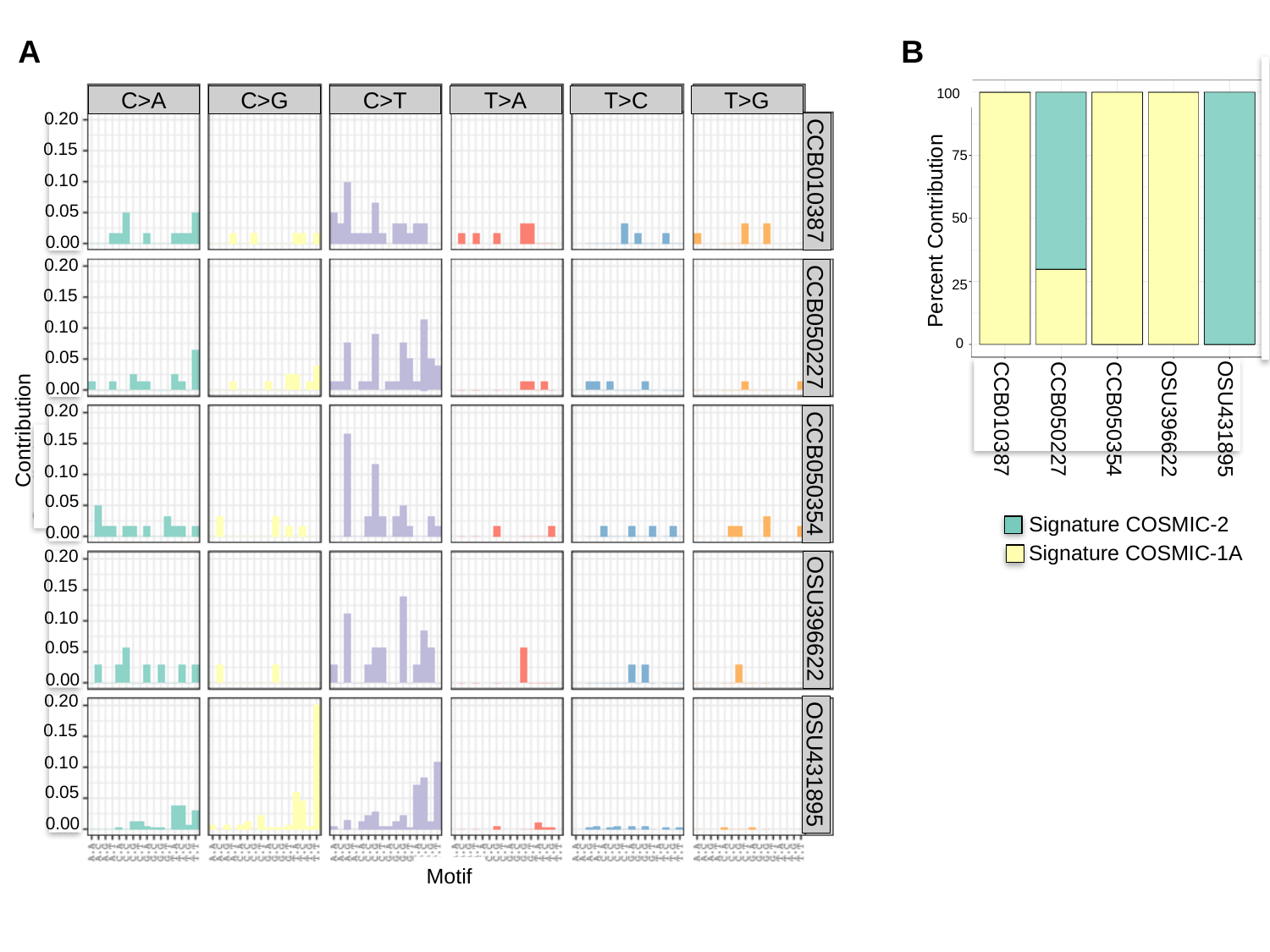

A
T>C
C>A
C>G
C>T
T>A
T>G
CCB010387
CCB050227
CCB050354
OSU396622
OSU431895
0.20
0.15
0.10
0.05
0.00
0.20
0.15
0.10
0.05
0.00
0.20
0.15
0.10
0.05
0.00
Contribution
0.20
0.15
0.10
0.05
0.00
0.20
0.15
0.10
0.05
0.00
Motif
B
100
75
50
Percent Contribution
25
0
CCB010387
CCB050227
CCB050354
OSU396622
OSU431895
Signature COSMIC-2
Signature COSMIC-1A
