## Supplemental Figure 4 for "Identification of frequent activating *HER2* mutations in primary canine pulmonary adenocarcinoma"

### Slide 1
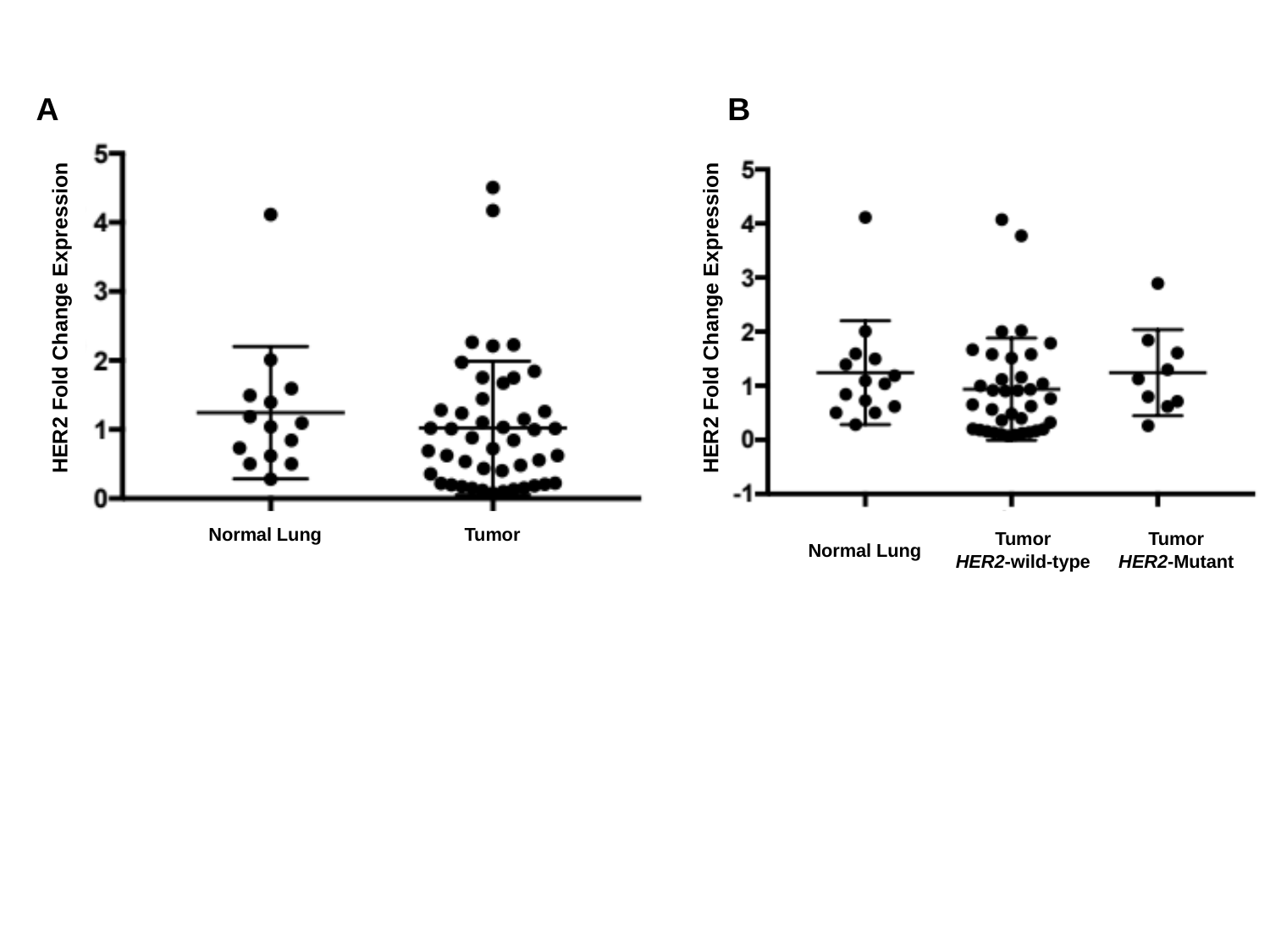

A
B
HER2 Fold Change Expression
HER2 Fold Change Expression
Normal Lung
Tumor
Tumor
HER2-wild-type
Tumor
HER2-Mutant
Normal Lung
